# Neural and computational processes of accelerated perceptual awareness and decisions: A 7T fMRI Study

**DOI:** 10.1101/2021.07.22.453451

**Authors:** Shivam Kalhan, Jessica McFadyen, Naotsugu Tsuchiya, Marta I. Garrido

**Affiliations:** School of Psychological Sciences, University of Melbourne, Melbourne VIC 3010 Australia; Max Planck UCL Centre for Computational Psychiatry and Ageing Research, University College London, London, United Kingdom; Queensland Brain Institute, University of Queensland, Brisbane, QLD 4072 Australia; School of Psychological Sciences, Faculty of Biomedical and Psychological Sciences, Monash University, Clayton, VIC, Australia; Monash Institute of Cognitive and Clinical Neuroscience, Monash University, Clayton, VIC, Australia; Australian Research Council Centre of Excellence for Integrative Brain Function

## Abstract

Rapidly detecting salient information in our environments is critical for survival. Visual processing in subcortical areas like the pulvinar and amygdala have been shown to facilitate unconscious processing of salient stimuli. It is unknown, however, if and how these areas might interact with cortical networks to facilitate faster conscious perception of salient stimuli. Here we investigated these neural processes using 7T functional Magnetic Resonance Imaging (fMRI) in concert with computational modelling while participants (n = 32) engaged in a breaking continuous flash suppression paradigm (bCFS) in which fearful and neutral faces are initially suppressed from conscious perception but then eventually “breakthrough” into awareness. We found that participants reported faster breakthrough times for fearful faces compared to neutral faces. Drift-diffusion modelling suggested that perceptual evidence was accumulated at a faster rate for fearful faces compared to neutral faces. For both neutral and fearful faces, faster response times coincided with greater activity in the amygdala (specifically within its subregions, including superficial, basolateral and amygdalo-striatal transition area) and the insula. Faster rates of evidence accumulation coincided with greater activity in frontoparietal regions and the occipital lobe, as well as the amygdala. Overall, our findings suggest that hastened perceptual awareness of salient stimuli recruits the amygdala and, more specifically, is driven by accelerated evidence accumulation in fronto-parietal and visual areas. In sum, we have uncovered and mapped distinct neural computations that accelerate perceptual awareness of visually suppressed faces.

## Introduction

At any given moment, we receive vast quantities of information about our ever-changing environments. To survive, it is critical that we can rapidly detect biologically-relevant stimuli within this stream of information. Previous studies have found that salient information (e.g. spiders, fearful faces, fear-conditioned stimuli) gain preferential access to awareness and are perceived faster than non-salient information (Gayet et al., 2016; Gomes et al., 2017). Faster detection of salient information might be facilitated by subconscious processing within subcortical neural networks (i.e. the ‘low-road’ subcortical route to the amygdala; Tamietto and de Gelder, (2010)). This ‘low-road’ describes a connection from the superior colliculus to the amygdala via the pulvinar that effectively bypasses cortical visual networks, thus enabling faster transmission to the amygdala (McFadyen et al., 2017) for processes relating to biological relevance and saliency computation (Koller et al., 2019).

An alternative hypothesis is that there are ‘many-roads’ for processing salient information, such that other neural networks (e.g., dorsal and ventral visual streams, as well as fronto-parietal and cingulate networks) play a significant role in the fast detection of salient information (Pessoa and Adolphs, 2010). Both hypotheses have in common that the amygdala plays a role in these processes. However, the hypotheses differ as to whether subcortical regions are also involved. The ‘many-roads’ hypothesis suggests a role of the amygdala in evaluating salience in concert with cortical networks. The ‘low-road’ hypothesis emphasises the role of the subcortical route to the amygdala with respect to rapid processing of survival critical stimuli.

The aim of the present study was to use functional magnetic resonance imaging (fMRI) to map neural activity in subcortical and cortical visual networks while participants made perceptual decisions about fearful and neutral faces. We used a breaking continuous flash suppression (bCFS) paradigm (Jiang et al., 2007; Stein et al., 2011, 2014), in which stimuli gradually emerged into conscious perception. Participants were tasked with reporting whether each stimulus was rotated clockwise or anticlockwise. The time taken to make this perceptual decision was used as a proxy for the time taken for the stimulus to enter peoples’ awareness.

First, we sought to establish whether different neural networks were recruited for neutral vs. fearful faces presented under bCFS. Previous accounts using the continuous flash suppression (CFS) paradigm (Tsuchiya and Koch, 2005) have used fMRI to establish differences between fearful and neutral stimuli when suppressed vs when consciously perceived (Jiang and He, 2006; Vizueta et al., 2012). Both studies found the involvement of the amygdala when processing suppressed fearful faces. Consistent with these studies, we hypothesised that the amygdala will be more strongly activated when processing fearful than neutral faces in the bCFS paradigm. Other studies, however, have found that activity in cortical regions (fronto-parietal, temporal, and occipital) correlates with perceptual transitions during binocular rivalry (Frässle et al., 2014; Knapen et al., 2011; Lumer et al., 1998; Lumer and Rees, 1999). Thus, we hypothesised that greater amygdala activity for fearful faces would coincide with increased activity within cortical networks, consistent with the ‘many-roads’ hypothesis (Pessoa and Adolphs, 2010). Interestingly, Balderston et al., (2017) used 3T-fMRI and tractography to show that there are functional differences between the distinct human amygdala subregions as they process fear stimuli. Therefore, there may also be intra-amygdala differences in the processing of suppressed fear and neutral stimuli. To test this possibility, we utilised a higher resolution 7T-fMRI where we were able to image specific amygdala subregions while participants completed our task.

Secondly, we manipulated expectations about emotional expression of the upcoming suppressed faces by using probabilistic cues. Previous research has found that more probable stimuli become consciously accessible earlier than improbable stimuli – i.e., “we see what we expect to see” (McFadyen et al., 2019; Melloni et al., 2011; Pinto et al., 2015; Sterzer et al., 2008). Thus, we sought to investigate whether the degree of surprise (or expectation) about a visual stimulus might modulate neural activity relating to neutral and fearful face processing during bCFS.

Finally, we utilised drift diffusion modelling (Ratcliff, 1978) to specifically address the computational mechanisms underlying differences in response times to neutral vs. fearful and expected vs. unexpected stimuli. Previous studies have found that the drift rate parameter (emuating the rate of evidence accumulation) is higher for biologically-relevant stimuli such as snakes and emotional facial expressions (Lerche et al., 2019; Lufityanto et al., 2016; Tipples, 2015). This increased rate of evidence accumulation (drift-rate) allows for faster accumulation of information and possibly leads to faster awareness of the stimulus and/or decision-output in response. To better understand the neural computations underlying faster response times, we also mapped neural activity to the drift diffusion model parameters. Overall, using the current task design, we were able to investigate the specific neural and computational processes involved in hastening perceptual awareness of suppressed visual stimuli.

## Methods

### Participants

We recruited 32 participants through the University of Queensland’s Participation Scheme. The sample consisted of 16 males and 16 females between the ages of 18 and 36 years (mean = 23.25 years, SD = 4.79). All participants reported having no colour-blindness and normal vision with or without corrective lenses. Participants with corrective glasses were excluded due to a practical difficulty with installing red-green anaglyph lenses on top of the prescription glasses in the MRI scanner. Participants with contact lenses were included in the present study. All participants were compensated AU$20 per hour for their time and provided written consent. This study was approved by the University of Queensland’s Human Research Ethics Committee.

### Stimuli

Face stimuli used in the present study were collected from different experimentally-validated databases, with the aim of maximising the number of unique faces presented. These databases included the Amsterdam Dynamic Facial Expressions Set (ADFES; van der Schalk et al., 2011), the Karolinska Directed Emotional Faces set (KDEF; Goeleven et al., 2008), the NimStim set (Tottenham et al., 2009) and the Waesaw Set of Emotional Expression Pictures (WSEFEP; Olszanowski et al., 2014). The overall set of face stimuli consisted of 267 images of Caucasian adults (66 females and 71 males), with either neutral or emotional facial expressions. We cropped the hair, neck and shoulders from each image and then centred the faces within a 365 × 365-pixel square with a black background. We calculated the luminance and root-mean square contrast of the entire image (consisting of the grey-scaled face and background), across all images using the SHINE toolbox (Willenbockel et al., 2010). There were no significant differences between the neutral and fearful emotional faces (luminance: neutral = 125.9, fearful = 124.68, t(130) = 1.954, p = 0.1; contrast: neutral = 125.9, fearful = 125.5, t(130) = 2.038, p = 0.09; Bonferroni-corrected for two comparisons).

To suppress the face stimuli from perceptual awareness, we used Mondrian images made using code available online (http://martin-hebart.de/webpages/code/stimuli.html; as also used in Stein et al., 2014). These Mondrian masks were presented at 125% the size of the face stimuli to ensure that the faces were sufficiently masked. We then used MATLAB to convert all face and mask stimuli into sets of either red or green by removing the non-relevant colour planes (e.g., to make red images, we removed the blue and green planes of the image). Having sets of red and green faces and masks allowed us to achieve dichoptic presentation with red-green anaglyph glasses (specific dimensions and materials of the lenses can be found here: http://www.oz3d.com.au/catalog/product/view/id/99/s/red-green-paper-3d-glasses/).

### Procedure

Participants first completed the MRI safety questionnaires and the consent form. We then determined the participants’ ocular dominance using the Miles Test (Miles, 1930). Participants wore the red-green anaglyph lenses such that the colour of the lens over the dominant eye matched the colour of the mask, which was predetermined using a counterbalancing approach to limit any confound of the red-green colours. Participants then completed a short practice task using a titration procedure (see *Behavioural Titration*). Participants were then moved into the MRI scanner, where they first performed a titration task to account for individual differences in differences in sensitivity to perceptual suppression caused by the mask and then proceeded with the main task (see *Behavioural Paradigm*). After completing scanning, participants completed the Beck’s Anxiety and Depression Inventory (21 questions each; Beck et al., 1996; Beck and Steer, 1990) outside of the scanner.

#### Behavioural Paradigm

For the behavioural task, we utilised a “breakthrough” variant of CFS (Jiang et al., 2007), which suppresses a stimulus of interest to one eye from conscious perception by presenting a high-contrast flickering mask to the other eye. In the bCFS paradigm, the contrast of the stimulus of interest is then gradually increased from low to high such that the stimulus is no longer suppressed by the mask and emerges into awareness. By incorporating a discrimination task for the stimulus of interest (here, whether the face was rotated clockwise or anticlockwise), the task response time measures when different types of stimuli became consciously discriminable (Stein et al., 2011).

In the present experiment, each trial began with a 0.7-second cue – the word “NEUTRAL” or “FEARFUL” – for the upcoming stimulus. The cues were probabilistic, such that they reliably predicted the upcoming emotional expression 80% of the time. After the cue, a 3-second transitioning anaglyph was presented. In the first frame, the mask stimulus (in the dominant-eye’s lens colour) was presented in the centre of the screen (see *Figure 1*). Over a period of 3 seconds, the face stimulus (in the non-dominant eye’s lens colour) was gradually superimposed over the mask stimulus at increasingly greater contrast levels (0 to 100% of titrated contrast level). The final frame of the trial consisted of both the mask and face simultaneously presented, both at 100% in different colours (red or green). On each trial, the face stimulus was rotated by 5° either clockwise or counter-clockwise in a pseudo-random order. Throughout the 3 seconds of each trial, the mask randomly changed to different Mondrian images at a rate of 10 Hz, creating a flickering effect that enhanced suppression of the face from conscious perception (Tsuchiya and Koch, 2005). A fixation cross was presented in the centre of the screen at all times. The inter-trial interval was uniformly and randomly jittered between 0.25 and 0.5 seconds in 0.05-second bins. Participants were instructed to report the rotation of the face as soon as it was perceptible by pressing the left (for counter clockwise) or right (for clockwise) button. Participants were instructed to be as fast as possible and were also informed that the emotional expression of the face was task-irrelevant.

**Figure 1.**
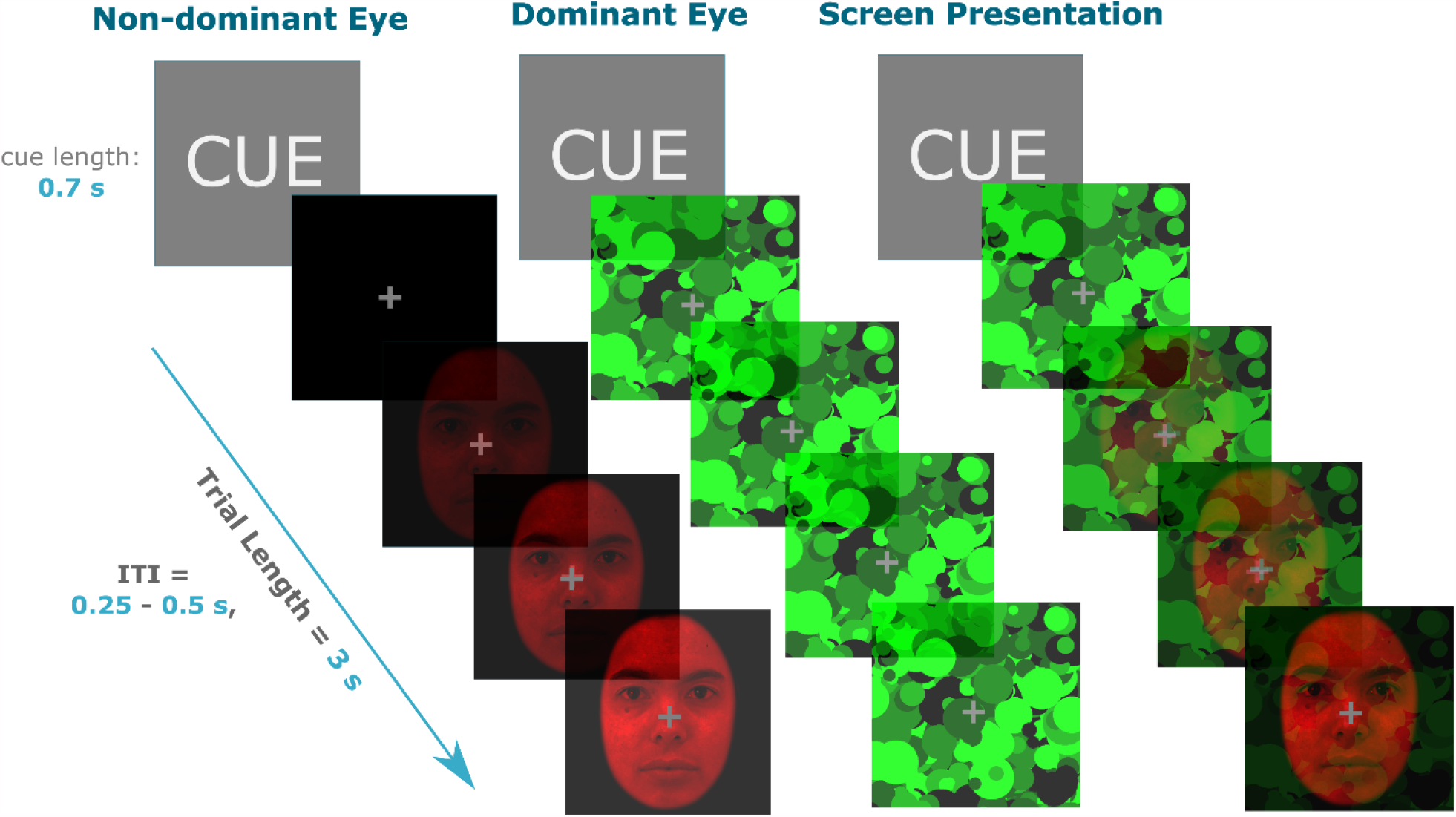
bCFS paradigm design example trial. Prior to the onset of the trial, participants were cued on the upcoming emotion of the face for 0.7s. The face and masks were then presented to each eye, with the face increasing in contrast for 3 seconds and mask at 100% contrast for the full duration. The screen presentation panel is what the trial looked like without the red-green anaglyph lenses, which were used to achieve dichoptic presentation. The face stimuli were always presented in the non-dominant eye and mask in the dominant eye. The inter-trial interval (ITI) was 0.25 – 0.50s.

Our design produced four different conditions: 1) expected fearful (EF; “fearful” cue, followed by a fearful face, 2) expected neutral (EN; “neutral” cue, followed by a neutral face), 3) unexpected fearful (UF; “neutral” cue, followed by a fearful face) and 4) unexpected neutral (UN; “fearful” cue, followed by a neutral face). There were 600 trials in total, consisting of 240 EF, 240 EN, 60 UN and 60 UF trials. These were all pseudo-randomly ordered with the following constraints: 1) no two unexpected trials presented in a row, and 2) no more than four consecutive presentations of the expected trials of the same emotional face. These 600 trials were split into 4 runs, containing 150 trials each. Each run had all four trial types with approximately 80-20% frequency of the expected-unexpected trials. The first five trials of each run were always expected trials. Lastly, the order that each run was presented was counter-balanced across participants to minimise any confounding order-effects. At the end of each run, participants had a less than 1-minute break while the scanner was restarted. The total task took approximately 42 minutes to complete (approximately10 minutes per run).

#### Behavioural Titration

A titration task was used to determine the relative contrast of the face and mask stimuli that produced average response times of approximately 2 seconds. This was to ensure that participants were able to respond within the 3-seconds trial period, while also accounting for individual differences in sensitivity to interocular suppression. The titration task contained all four conditions (EN, EF, UN, UF) as presented in the main behavioural task.

The first trial in the procedure started with the mask at lowest contrast relative to the face at highest contrast (100% face, 0% mask). We then altered their contrast using a staircasing procedure. Specifically, if the response time was faster than 2 s, the contrast of the mask was increased on the subsequent trial while that of the face was decreased so that the sum of the contrasts was always 1. Conversely, if response time was slower than 2 s, the contrast of the face on the following trial was increased (and the mask contrast decreased). This was adjusted trial-by-trial using the Palamedes toolbox in MATLAB (Prins and Kingdom, 2009) using a stepwise function, starting with 10% contrast adjustments that were then decreased by 2% after each reversal (i.e., a change in response type; fast to slow, or slow to fast). After four reversals, contrast adjustments were fixed at 2%. The procedure produced four sets of contrast values per condition. A single contrast value was then calculated by taking the mean contrast across all conditions (EN, EF, UN, EF), which was subsequently used for all trials in the main task. The mean response time from the main task across all trials and participants was 1.9 s; a close match to the titrated value of 2 s.

Participants performed a brief (5 minutes, 80 trials) practice titration task before entering the scanner to ensure participants understood the task. This practice task did not include the unexpected conditions. The full titration procedure (120 trials) then took place inside the scanner while structural T1 scans were acquired.

#### MRI Image Acquisition Parameters and Sequences

Imaging data was acquired using a MAGNETOM 7T whole-body scanner (Siemens Healthcare, Erlangen, Germany) with a 32-channel head coil (Nova Medical, Wilmington, US). The parameters for echo-planer imaging (EPI) sequences for whole-brain coverage used were the following: TR = 1.58s, TE = 21ms, flip angle = 65°, voxel size = 1.5 × 1.5 × 1.25mm^3^, matrix dimensions = 148 × 148 × 78. The sequences were chosen based on Sladky et al., (2018), optimised for imaging the amygdala.

### Analysis

#### Behaviour

Trials that had a response time of 0.5 s or less were removed. We also removed outliers (outside 3 standard deviations). To determine response time differences between conditions we used a 2 (prediction: expected, unexpected) x 2 (emotion: neutral, fearful) repeated-measures analysis of variance (ANOVA) on only correct trials (mean accuracy = 94%).

#### Drift Diffusion Modelling

We used drift diffusion modelling (DDM) (Ratcliff, 1978) and estimated its parameters to explain the response time distribution. We then used the estimated parameters to localise the respective neural correlates of psychological processes that are captured by these parameters. For parameter optimisation we tested a series of eight models, that is, a power set (=8) of three parameters (decision-boundary: α, drift rate: v, and non-decision time: t0) per condition. To test for which model best explained our data, we used the *fast-dm* software that estimates maximum likelihood (Voss and Voss, 2007). Other than α, v, and t0, we fixed three available parameters based on our experimental design: 1) starting point (*z*) at 0.5 as the face rotation was randomised, differences in response execution at 0 in order to reduce the number of free parameters and we expect that the decision boundary has the same effect for adjusting response execution in our task, inter-trial variability was fixed at 0 for *z* and *v* as the number of incorrect trials (average of 5% across all conditions) were not enough to give reliable inter-trial estimates. We then computed the Akaike information criteria (AIC) and the Bayesian information criteria (BIC) individually for each participant (all conditions) to test for which of the 8 models best explained our data at the group level (by summing AIC/BIC across participants for each model). We then extracted the parameter estimates from the winning model for each of the four conditions for each participant. We compared the differences between these condition-specific parameter estimates using a 2 (prediction: expected, unexpected) x 2 (emotion: neutral, fearful) ANOVA design with the two factors.

#### Functional Magnetic Resonance Imaging

We recruited SPM12 (http://www.fil.ion.ucl.ac.uk/spm/software/spm12/) for all imaging analyses. We pre-processed the MRI data using SPM default functions with steps in the order of realignment, co-registration, and segmentation, and normalised into MNI-space with spatial resolution of 1.5 × 1.5 × 1.5 mm^3^. We smoothed the images using a 6mm FWHM Gaussian kernel. We excluded runs with movement of greater than 3 mm (double the voxel size) in either of the x, y, z planes from further analyses. All first-level general linear models (GLMs) included a regressor for the four conditions (EF, EN, UF, UN) as well as six additional movement regressors. We applied a 128 s high-pass filter and set the masking threshold to 0.4, specified in the first-level modelling. We performed all functional analyses within the time epoch of 1.9 s (mean response time across all conditions and participants) from the onset of the face stimuli. We designed a total of three second-level GLMs that included the four condition regressors, as well as either: GLM 1) no additional regressor, GLM 2) average response time and, GLM 3) drift rate. All these were added in the second-level modelling, with any response-time and drift-rate outliers (+/- 3 standard deviations) removed. We determined the anatomical labels for the resultant maps of significant neural activity using the SPM Anatomy Toolbox (Eickhoff et al., 2005).

We also conducted ROI analyses to identify activation within the amygdala and its subregions that covaried with response time, drift-rate and decision-boundary using the 4 GLMs described above. We created an amygdala-specific anatomical mask (using SPM Anatomy Toolbox; Eickhoff et al., 2005) by combining individual masks of four of the amygdala subregions: superficial (SF), amygdalo-striatal transition area (AStr), basolateral (BL) and centromedial (CM). This mask was specified in the second-level analysis as an explicit mask for the ROI analysis. The background image in all brain figures is from Bollmann et al., (2017); https://imaging.org.au/Human7T/MP2RAGE). All fMRI data presented is at p < 0.05 family-wise error corrected threshold, with a voxel cut-off of k = 4.

## Results

### Behaviour

#### Perceptual decisions are accelerated for fearful faces

We first investigated the response time data across conditions. Both the fearful conditions (EF; 1.87s ± 0.03 and UF; 1.88s ± 0.03) had a faster breakthrough time compared to the neutral conditions (EN; 1.91s ± 0.03 and UN; 1.92s ± 0.03) - main effect of emotion: F(1,30) = 21.56, p < 0.001. There were no significant response time differences based on the prediction factor (main effect of prediction: F(1,30) = 2.2, p = 0.15) and no interaction between the emotion and prediction factors (F(1,30) = 0.56, p = 0.46). Overall, this suggests that both fearful conditions (EF and UF) had a faster response time compared to the neutral conditions (EN and UN), irrespective of expectation (see Figure 2). Accuracy scores were at ceiling across all four conditions: EN (95% ± 0.8), UN (95% ± 0.7), EF (94% ± 0.9) and UF (96% ± 0.6).

**Figure 2.**
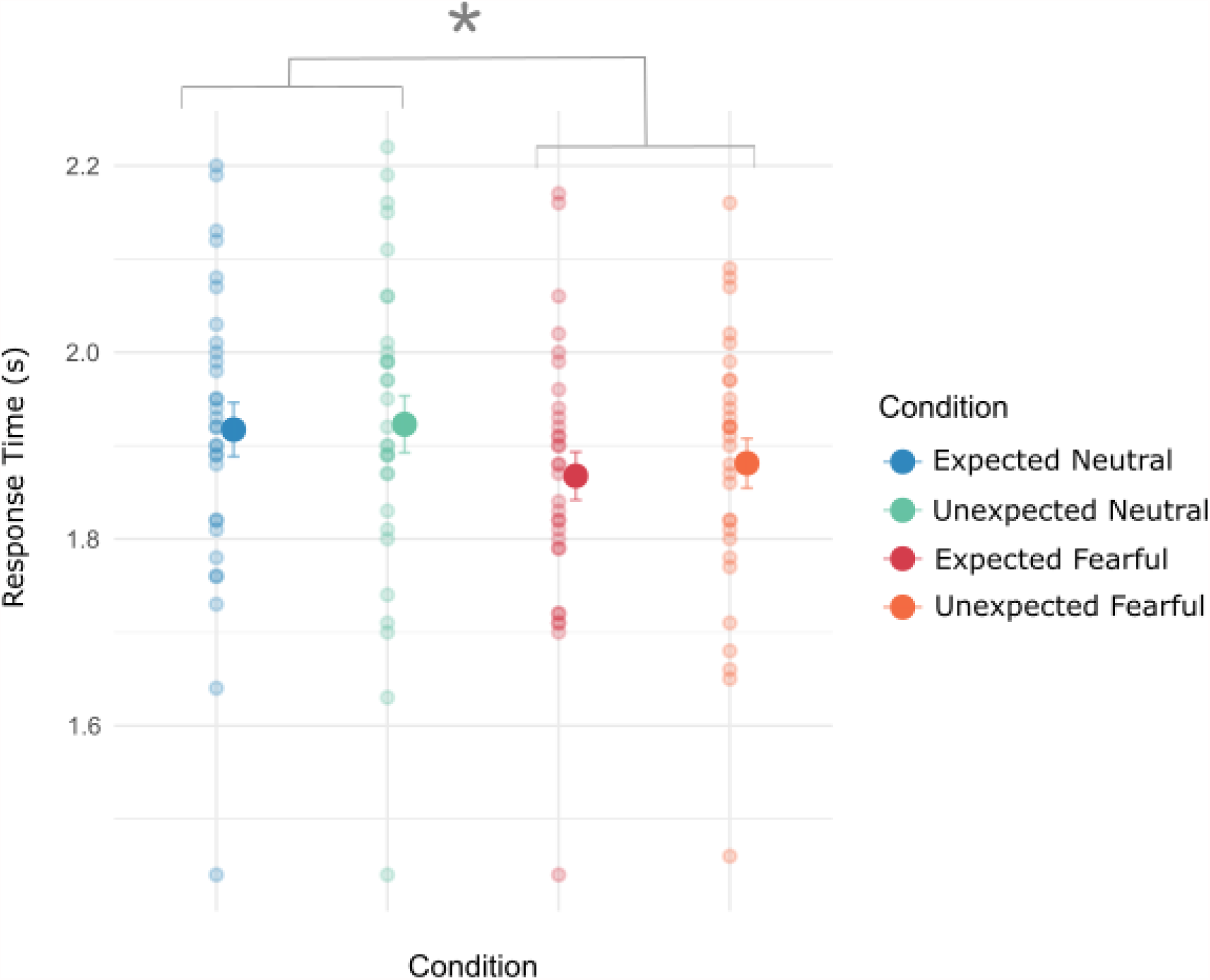
Behavioural results of response time and accuracy across conditions. Fearful faces (both EF and UF) had faster response times compared to neutral faces. Smaller dots are individual participants’ mean response times, with the larger dot being the overall mean. *\* p <* .*05*

#### Faster drift-rate for fearful faces

We used drift-diffusion modelling to investigate the processes that underpin accelerated response time we observed for fearful faces. We first performed parameter optimisation to determine the winning model (see Methods for details). To determine this, we computed the AIC and BIC scores and compared across the eight models. We found that model 3, which included the free parameter of drift-rate (v), best explained our data according to AIC and BIC (by 3.5 over the second-best model, *m2*) (see *Figure 3*). Therefore, we decided to report the condition specific differences of these parameters across participants using model 3 (*v*).

**Figure 3.**
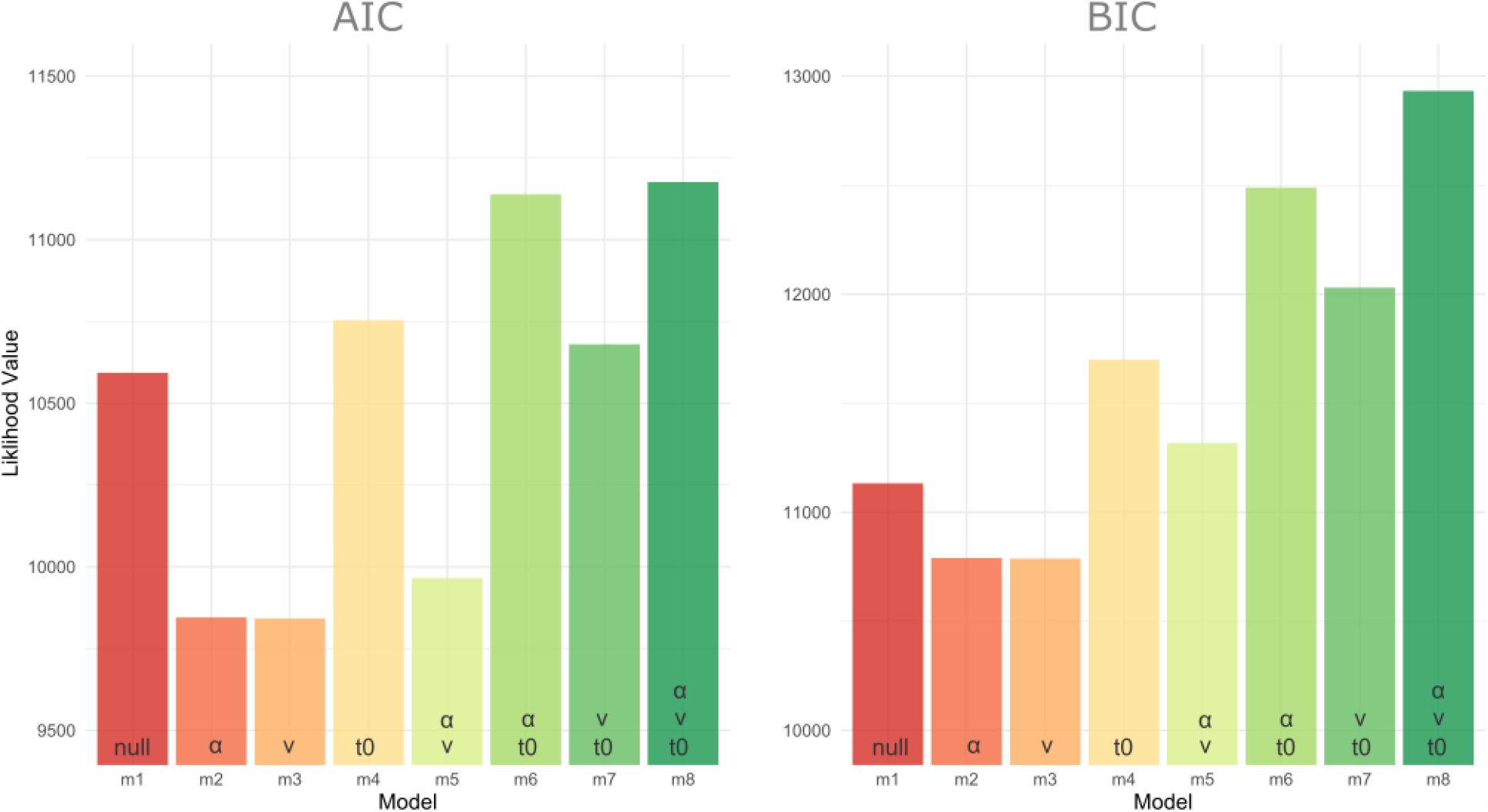
AIC and BIC scores for parameter optimisation across all eight models (m_1-8_). Parameter combinations used for each model are specified at the bottom of the bars. Lower values indicate better fits. For both AIC and BIC, the winning model was model 3 (*v*), by 3.5, compared to the next best model (model 2; *α). v = drift-rate, α = decision-boundary, t*_*0*_ = *non-decision time*.

Fearful faces (both expected and unexpected) had a significantly faster drift-rate (main effect of emotion; F(1,27) = 20.12, p < 0.0001) compared to neutral faces (EN = 1.66 ± 0.06, UN = 1.65 ± 0.05, EF = 1.74 ± 0.06, UF = 1.72 ± 0.06). We did not find a significant main effect of prediction (F(1,27) = 0.52, p = 0.5), nor an interaction between emotion and prediction factors (F(1,27) = 0.1, p = 0.75). This suggests that faster response times for the fearful conditions were influenced by an increased drift-rate (*Figure 4*), which reflects an increased rate of evidence accumulation of the fearful stimuli and facilitate a faster perceptual discrimination on the rotation of the faces.

**Figure 4.**
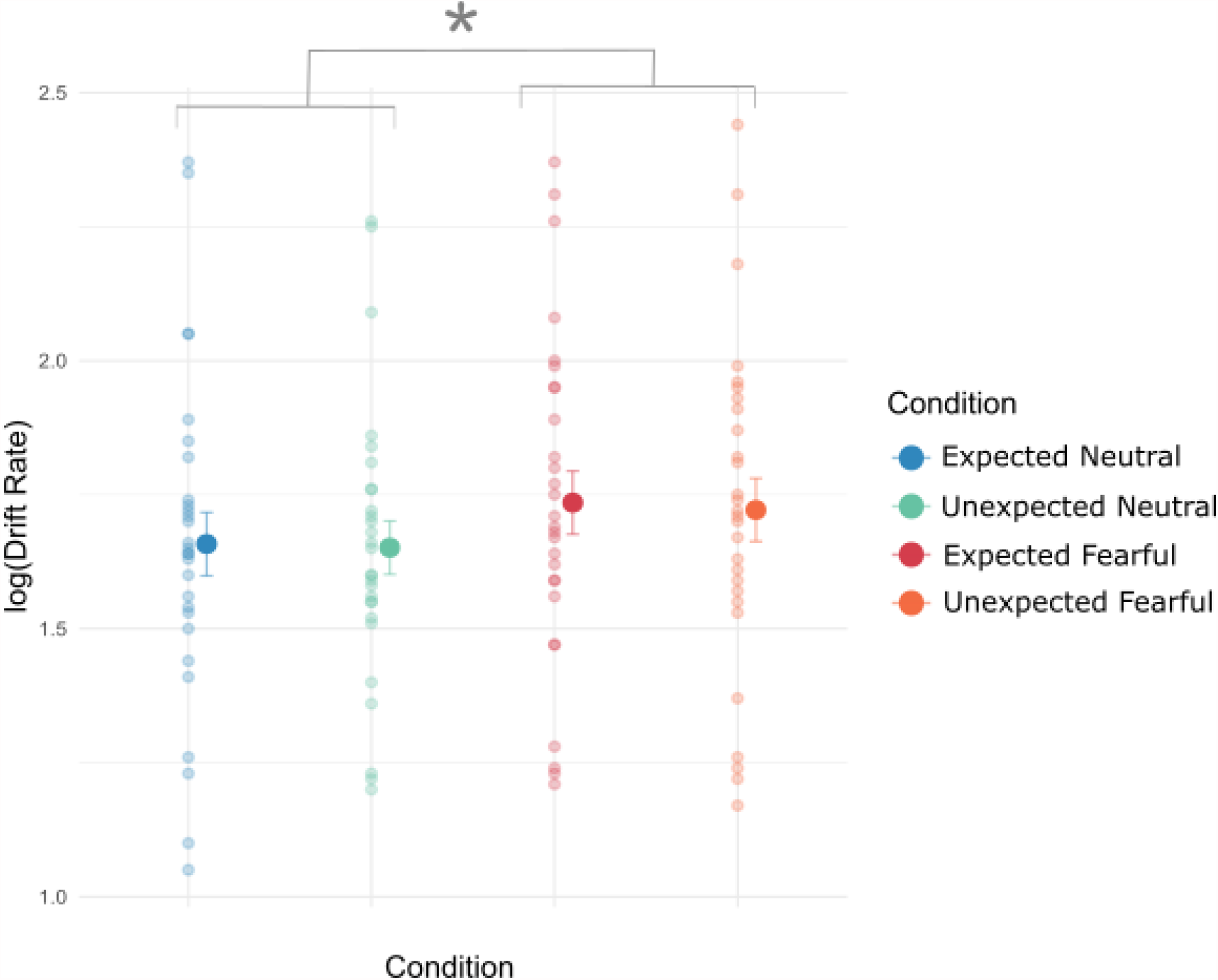
Parameter estimates across conditions using drift-diffusion modelling. Drift-rate parameter estimates. Fearful faces had a faster drift rate compared to neutral faces. Individual small dots represent each participant’s parameter estimate. The mean is the black dot inside the density plot. *Results plotted as mean ± SE. * p <* .*05*

#### fMRI Neural Activity

##### Neural correlates of accelerated response times

We first investigated patterns of whole-brain BOLD signal that correlated with faster response time (negative T-contrast) across all conditions (GLM 2). We found significant covariation in the bilateral insular lobes, bilateral Rolandic operculum which covers the insula (top panel, Figure 5), bilateral cuneus (bottom left, Figure 5), the right visual thalamus (bottom right Figure 5), superior and middle temporal gyrus (MTG and STG), and the amygdala. We then performed an ROI analysis on the amygdala and its subregions (superficial (SF), amygdalo-striatal transition area (AStr), basolateral (BL) and centromedial (CM)). There was a small cluster (9 voxels) in the left amygdala within the SF area that significantly covaried with response time. In the right amygdala, we observed a larger cluster (117 voxels) across the AStr, SF and BL areas. A large percentage of AStr (82.3%) was activated, with 25.9% of this cluster being within the AStr. We also found a slightly larger percentage of this cluster to be within the SF area (31.7%), with 32.5% of this subregion activated. Lastly, 25.6% of this cluster was also within the BL with 5.9% percentage of this area activated (Figure 6; see supplementary material (Table S1) for a list of all the brain regions activated and their respective MNI-coordinates).

**Figure 5.**
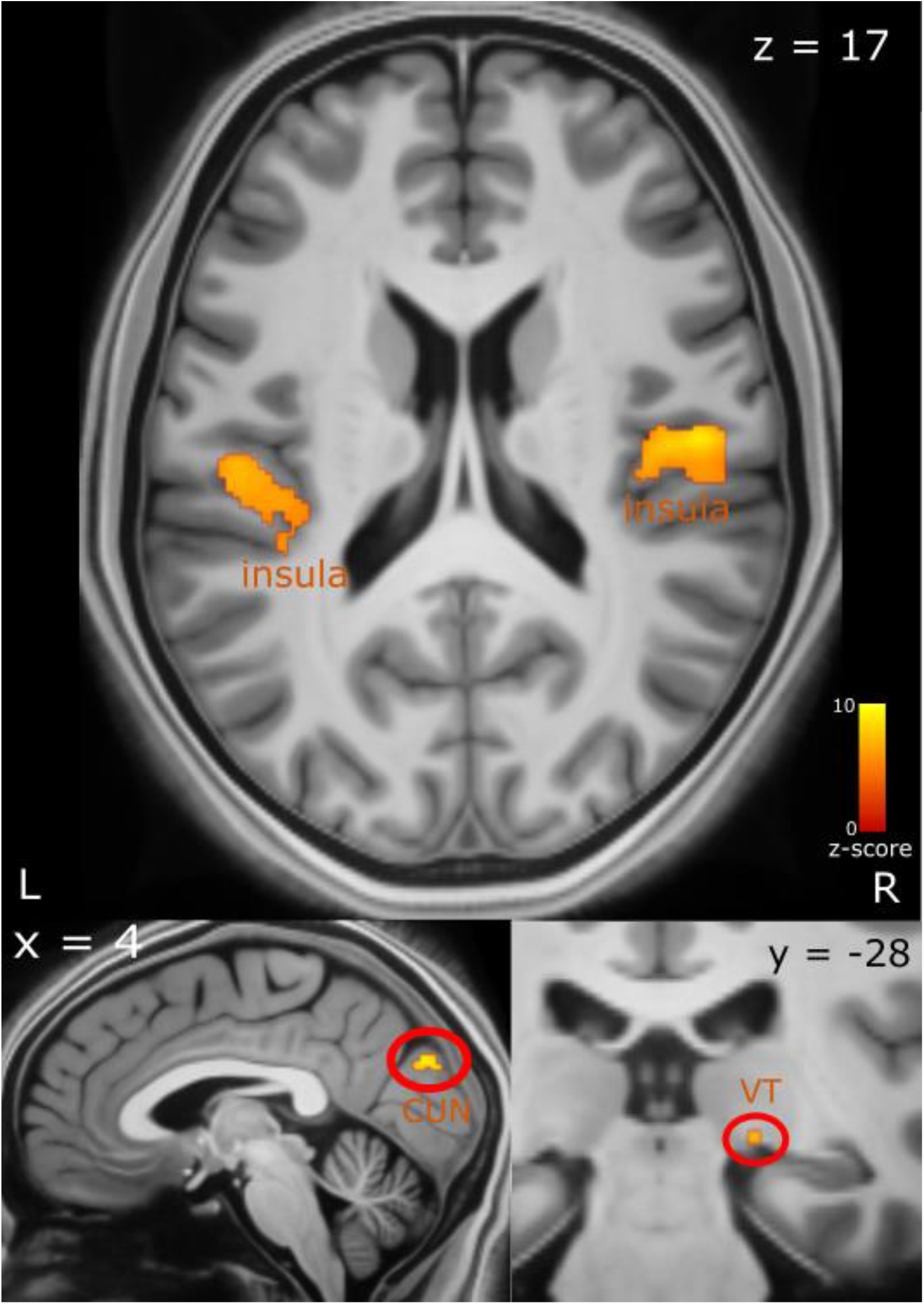
Brain activity correlated with faster response times. Top panel shows activation in the insula and Rolandic Operculum. The bottom left panel shows activation in the cuneus and bottom right in the right visual thalamus. The axial (z), coronal (y) and sagittal (x) MNI coordinates are embedded in the relevant images. *L = left, R = right. CUN = cuneus, VT = visual thalamus*.

**Figure 6.**
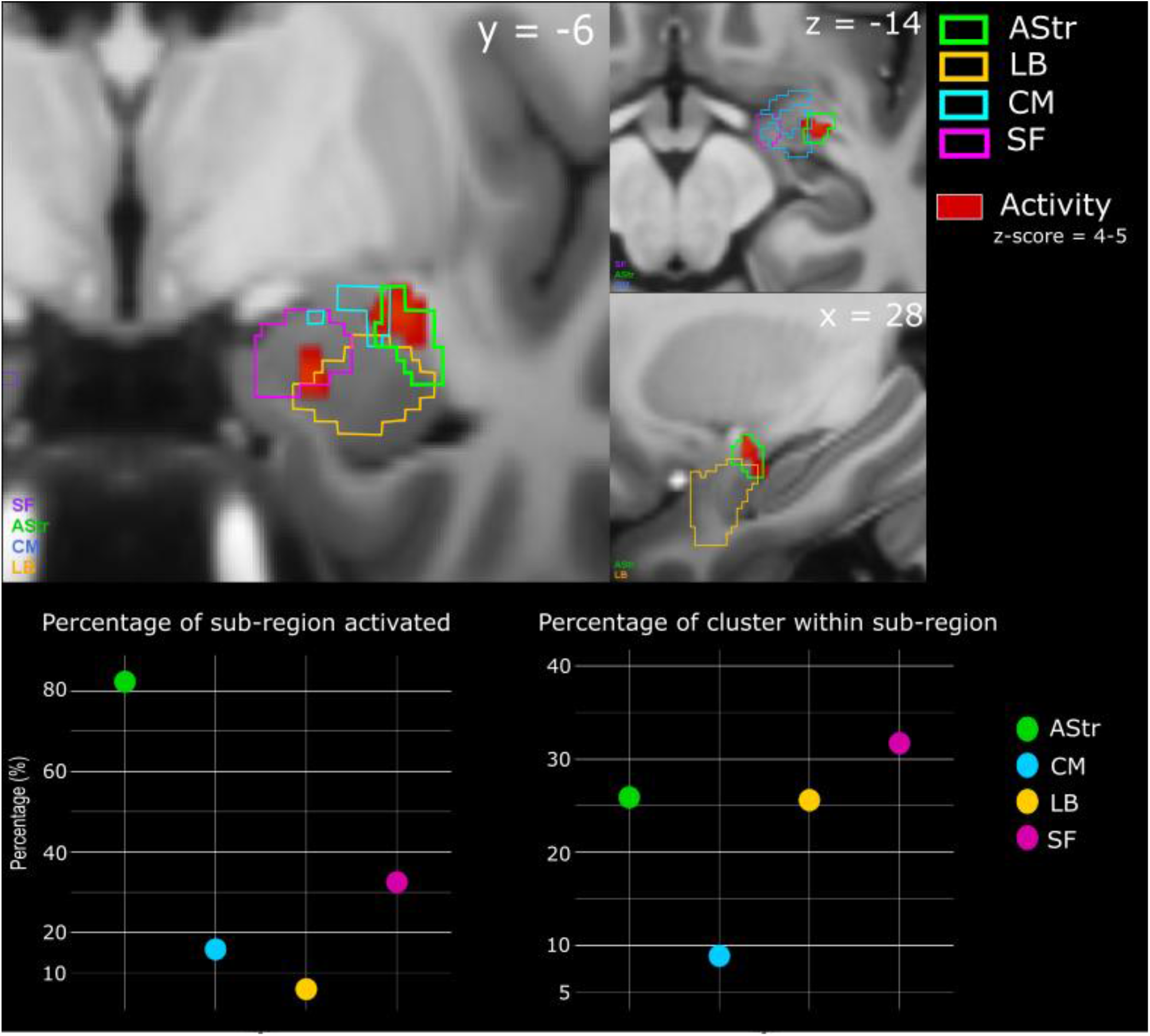
Subregion activations within the right amygdala, correlated with faster response time. Top panel shows activity in red with the subregional divisions overlaid in their respective colours. The bottom left graph shows the percentage of subregion activated. The bottom right graph shows the percentage of the cluster within each of the subregions. The axial (z), coronal (y) and sagittal (x) MNI coordinates are embedded in the relevant images. *AStr = amygdalo-striatal transition area, SF = superficial, BL = basolateral, CM = centromedial*.

##### Neural correlates of increased drift rate/evidence accumulation

We next investigated which brain regions were engaged in the implementation of the computation modelled above, namely the increased drift-rate (positive T-contrast; GLM 3) or evidence accumulation. We found that greater activity in frontal regions including the middle frontal gyrus (MFG) and inferior frontal gyrus (IFG) significantly covaried with increased drift rate (see Figure 7).

**Figure 7.**
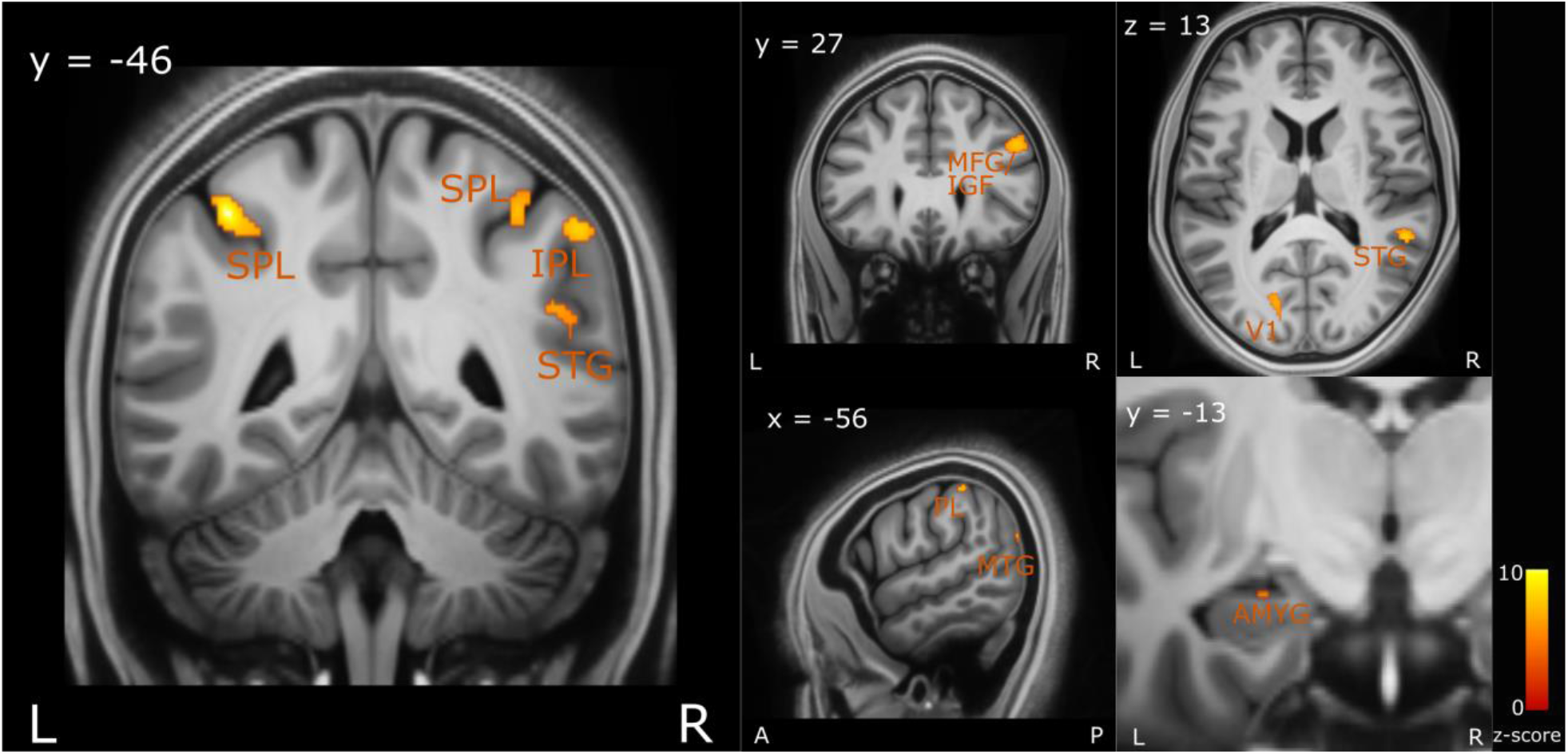
Brain regions correlated with increased drift-rate. The axial (z), coronal (y) and sagittal (x) MNI coordinates and brain region labels are embedded in the relevant images. *MFG = middle frontal cortex, MTG = middle temporal gyrus, V1 = primary visual cortex, IPL = inferior parietal lobule, PL = parietal lobe, SPL = superior parietal lobe, STG = superior temporal gyrus, AMYG = amygdala. L = left, R = right*.

There was also significant covariation within the primary visual cortex and the surrounding visual areas (including the middle and superior occipital gyri), as well as the middle and superior temporal gyrus and bilateral inferior and superior parietal lobe (IPL and SPL). The parietal lobe has been associated with attention, evidence accumulation and perceptual decision-making which are processes that are likely involved in accelerating the response time in our task (Behrmann et al., 2004; d’Acremont et al., 2013; Ploran et al., 2007). Lastly, there was also a relatively small cluster (voxels = 13) activated within the left amygdala. See supplementary material (Table 2) for a list of all the brain regions activated and their respective MNI coordinates.

## Discussion

Our findings demonstrate that participants can perceptually discriminate fearful faces faster than neutral faces in a breaking continuous flash suppression paradigm, replicating a previous finding from our lab (McFadyen et al., 2019) and supporting findings from previous psychophysical experiments (Jiang et al., 2009; Yang et al., 2007). Drift diffusion modelling revealed that the faster response times to fearful faces could be explained by an increased rate of evidence accumulation prior to response. Using 7T-fMRI we found that faster response times correlated with greater activity in the insula, amygdala, visual areas, and the temporal gyrus, across both fearful and neutral faces. Further, an increased drift rate correlated with activity in the parietal lobe, inferior and medial frontal gyri, as well as the temporal gyrus and amygdala, across both neutral and fearful faces.

The response time findings support previous literature in that fearful faces do gain preferential access to awareness. The faster perceptual discrimination and response times for fearful faces are computationally explained by a greater rate of evidence accumulation (drift-rate) as suggested by our drift-diffusion modelling results. These response time differences may therefore be influenced by fearful faces breaking into awareness faster than neutral faces, and also by fearful faces leading to faster perceptual decision-making in discriminating the face’s rotation. Our computational model suggested an involvement of these processes where fear hastened perceptual decision-making enabled by an increased drift-rate, allowing for a faster accumulation of visual evidence.

Contrary to our hypothesis, we did not observe significant brain activity differences between the fearful and neutral conditions (supplementary material, Figure S1). This is likely due to neutral faces having an equal relevance to fearful faces in completing our task, where participants were tasked to discriminate facial orientation irrespective of emotion. This interpretation is consistent with Reinders et al., (2005), where both fearful and neutral faces were shown to engage salience related brain regions such as the amygdala, when contrasted with non-salient stimuli (e.g. a house). Additional studies have also demonstrated that neutral faces have a relatively high salience value, evident from the observed activation of salience-related brain regions in response to these neutral faces (Fischer et al., 2003; Goossens et al., 2009; Jiang and He, 2006; Kesler-West et al., 2001; Ottaviani et al., 2012; Reinders et al., 2006; Santos et al., 2011). Consistent with these previous accounts, we found that faster response times correlated with activity in the insula and amygdala across both, fearful and neutral conditions. The insula has been implicated in facilitating attentional processing, both externally towards the stimuli and internally with interoceptive processing. Moreover, the insula mediates visual awareness in CFS and binocular rivalry paradigms (Frässle et al., 2014; Knapen et al., 2011; Lumer et al., 1998; Lumer and Rees, 1999; Menon and Uddin, 2010; Salomon et al., 2018, 2016). Additionally, our findings support the theoretical proposal by Craig (2009) in that the insula plays a critical role in conscious awareness and salient information processing. Overall, insula activity correlated with faster response times and suggest that faster breakthrough and/or perceptual decisions are mediated by rapid salience detection, enhanced for both neutral and fearful faces.

The amygdala is a key component in salience processing (Ledoux, 1998; Pessoa and Adolphs, 2010), and is thought to play a role in general relevance/salience detection (Attar et al., 2010; Sander et al., 2003), including responding to faces irrespective of emotional valence (Goossens et al., 2009; Lin et al., 2020; Santos et al., 2011). The amygdala has also been shown to process visually-suppressed salient stimuli in CFS and binocular rivalry paradigms (Jiang and He, 2006; Pasley et al., 2004; Troiani and Schultz, 2013; Vizueta et al., 2012; Williams et al., 2004). Consistent with these previous accounts, we interpret amygdala activation in our data as playing a role in detecting stimulus salience and potentially increasing attention, thereby facilitating perceptual awareness and/or decision, for both neutral and fearful faces. More specifically, we show that the AStr subregion within the right amygdala plays a significant role in these processes, together with the BL and SF subregions. The rat AStr was shown to conduct given neural stimulation with high velocity *in vitro*, relative to other amygdala subregions and this was interpreted as AStr being involved in producing fast behavioural responses (Wang et al., 2002). Therefore, our finding of AStr’s involvement with hastening response times is consistent with the interpretation from Wang et al., (2002). However, the activation of the other subregions (SF and BL) within the amygdala also implicates that several different processes within the amygdala may be involved in hastening response times. For example, the BL of the amygdala receives sensory input (Davis and Shi, 2000), suggesting that visual processes and computations involved in fastening reaction times may be happening at a relatively early stage of visual processing. Additionally, the SF regions of the amygdala (compared to the deeper amygdala regions) had been shown to be better connected to the frontal cortex (Bach et al., 2011), and therefore the activation of SF as well as the frontal cortex (see Table S1 in supplementary materials) in our task may also suggest the involvement of SF-frontal cortex circuitry in hastening the response times. Goossens et al., (2009) combined 3T fMRI with cytoarchitectonic probability maps of the amygdala and found that the SF amygdala is generally activated in response to faces (with fearful, happy and neutral expressions) and these responses did not differ across different emotional expressions. Our finding, with a higher spatial resolution of 7T fMRI, is consistent with that of Goossens et al., (2009) since we also found SF activation across both fearful and neutral faces. Critically, the lack of intra-amygdala differences in processing fearful and neutral faces, supports the general salience processing hypothesis of the amygdala proposed by Sander et al., (2003). Lastly, we also found activation of temporal, occipital and frontoparietal cortical regions correlated with increased response times. This finding, together with insula activation, is consistent with the ‘many-roads’ hypothesis where there is involvement of various brain regions/pathways along with the amygdala in processing visually salient information (Pessoa and Adolphs, 2010). In sum, faster response times for both neutral and fearful faces appeared to engage the salience processing systems, with contributions from the insula and the amygdala.

Correlating our computational model parameters with the neuroimaging data across all conditions allowed for a more mechanistic understanding of the processes involved leading up to the perceptual awareness and decision, as well as where in the brain these processes might be implemented. Faster drift rate correlated with increased activation of the IFG and MFG, both part of the dorsolateral prefrontal cortex (dlPFC) (Cieslik et al., 2013). These regions have been shown to play a critical role in accumulating sensory evidence (Pleger et al., 2006; Summerfield et al., 2006), collectively facilitating perceptual decision-making by integrating outputs from lower-level task-related sensory regions (Heekeren et al., 2008, 2004). Consistent with previous work, our results show that dlPFC activity also co-occurs with activation in low-level visuo-sensory areas including the occipital lobe and MTG. Therefore, the dlPFC likely integrates visuo-sensory evidence from the occipital lobe and MTG, which allows for the accelerated decision-output in discriminating the face’s rotation. Further support for this comes from Philiastides et al., (2011). Philiastides and colleagues reported that the disruption of dlPFC using repetitive transcranial magnetic stimulation (rTMS) in a perceptual decision-making task slowed response times, which was associated with a reduced drift-rate. However, it is important to note that there have been inconsistent results using rTMS on dlPFC in assessing conscious awareness (see Bor et al., (2017) and Rounis et al., (2010)). However, overall, there is evidence demonstrating the integrative role of dlPFC in evidence accumulation and perceptual decision making.

Prior work using fMRI and the CFS paradigm found that the superior temporal sulcus (STS is involved in processing visible and suppressed fearful faces but not suppressed neutral faces (Jiang and He, 2006). Vizueta et al., (2012) also used the CFS paradigm in fMRI and found that the STS activity correlated with processing suppressed fearful faces, but only when accounting for negative affectivity traits. Critically, in both tasks, STS activity was positively correlated (across subjects) with activity in the amygdala. In contrast, in our task (using bCFS) we found activity in the MTG and the STG (which is anatomically separated by the STS) to be correlated with faster response time and increased drift rate, across both, neutral and fearful faces. Consistent with the previous studies, we also found amygdala activity (although a relatively small cluster for drift-rate correlation) co-occurring with regions close to the STS (the MTG and STG). However, contrary to the previous accounts, we also found MTG and STG/amygdala activity for neutral faces. Given that our behavioural task places equal demands/task relevance on both face conditions, it is likely that the MTG and STG/amygdala activity here is playing a role in processing both neutral and fearful faces. The contrasting findings may also suggest that different brain and computational processes are engaged in the bCFS task compared to the CFS paradigm, with bCFS likely engaging brain regions involved perceptual in decision-making, in addition to just perceptual awareness.

A limitation of the present study is that we were unable to disambiguate the specific contributions of the two processes of perceptual awareness and perceptual decision-making. This is due to the possibility of our response time capturing both the time taken to perceive the stimuli *and* the time taken to decide (discriminate rotation) and make a response. In the future, to better disambiguate the specific contributions of threat processing from the effects of low-level visual features inherent to fearful vs neutral faces, a controlled fear conditioning paradigm may allow for more explicit interpretations. Further, given that fearful faces have larger eye whites discriminating their orientation may be easier compared to neutral faces. This could therefore lead to the faster response times for fearful faces and may be a potential confound in the present study. Future studies may better account for this by using a localization or detection paradigm (del Río et al., 2018; Pinto et al., 2015). Lastly, contrary to our hypothesis, we did not find response time differences relating to the expectation manipulation. One explanation for this lack of an effect may be due to the task design where expectation had a relatively low task relevance. Participants were tasked to discriminate face rotation *irrespective* of the cue and therefore may have ignored the cue altogether. For this reason, the expectation factor may not have been encoded in the first place.

In sum, we find speeded response time in our task that correlated with salience-related brain regions for both neutral and fearful faces. In search of a more mechanistic account, we correlated our computational parameter estimate of drift rate with our brain data. We found that increased drift rate engaged regions within the dlPFC (IFG and MFG) as well as visuo-sensory (occipital lobe and MTG) and attentional brain regions (IPL), suggesting that collectively, these regions contribute to increased rate of sensory evidence accumulation that leads to faster perceptual awareness and decision processes. We also found that faster response time correlated with increased activity in the amygdala and the insula, likely playing a role in salience detection across both neutral and fearful faces. Overall, we shed light on the specific neural computational processes leading to awareness and perceptual decisions of salient information processing that as it emerges into consciousness.

## Acknowledgements

This work was funded by the Australian Research Council Centre of Excellence for Integrative Brain Function (ARC Centre Grant CE140100007) to M.I.G. We thank the participants for their time.

## References

Attar CH, Müller MM, Andersen SK, Büchel C, Rose M. 2010. Emotional Processing in a Salient Motion Context: Integration of Motion and Emotion in Both V5/hMT+ and the Amygdala. J Neurosci 30:5204–5210. doi:10.1523/JNEUROSCI.5029-09.2010

Bach DR, Behrens TE, Garrido L, Weiskopf N, Dolan RJ. 2011. Deep and Superficial Amygdala Nuclei Projections Revealed In Vivo by Probabilistic Tractography. J Neurosci 31:618–623. doi:10.1523/JNEUROSCI.2744-10.2011

Balderston NL, Hale E, Hsiung A, Torrisi S, Holroyd T, Carver FW, Coppola R, Ernst M, Grillon C. 2017. Threat of shock increases excitability and connectivity of the intraparietal sulcus. Elife 6:e23608. doi:10.7554/eLife.23608

Beck AT, Steer RA. 1990. Manual for the Beck Anxiety Inventory. Behav Res Ther.

Beck AT, Steer RA, Brown GK. 1996. Manual for the Beck depression inventory-II. San Antonio, TX Psychol Corp.

Behrmann M, Geng JJ, Shomstein S. 2004. Parietal cortex and attention. Curr Opin Neurobiol 14:212–217. doi:10.1016/j.conb.2004.03.012

Bollmann S, Janke A, Marstaller L, Reutens D, O’Brien K, Barth M. 2017. MP2RAGE T1-weighted average 7T model. doi:https://doi.org/10.14264/uql.2017.266

Bor D, Schwartzman DJ, Barrett AB, Seth AK. 2017. Theta-burst transcranial magnetic stimulation to the prefrontal or parietal cortex does not impair metacognitive visual awareness. PLoS One 12:e0171793. doi:10.1371/journal.pone.0171793

Cieslik EC, Zilles K, Caspers S, Roski C, Kellermann TS, Jakobs O, Langner R, Laird AR, Fox PT, Eickhoff SB. 2013. Is there one DLPFC in cognitive action control? Evidence for heterogeneity from Co-activation-based parcellation. Cereb Cortex 23:2677–2689. doi:10.1093/cercor/bhs256

Craig AD. 2009. How do you feel - now? The anterior insula and human awareness. Nat Rev Neurosci. doi:10.1038/nrn2555

d’Acremont M, Fornari E, Bossaerts P. 2013. Activity in Inferior Parietal and Medial Prefrontal Cortex Signals the Accumulation of Evidence in a Probability Learning Task. PLoS Comput Biol 9:e1002895. doi:10.1371/journal.pcbi.1002895

Davis M, Shi C. 2000. The amygdala. Curr Biol 10:R131. doi:10.1016/s0960-9822(00)00345-6

del Río M, Greenlee MW, Volberg G. 2018. Neural dynamics of breaking continuous flash suppression. Neuroimage 176:277–289. doi:10.1016/j.neuroimage.2018.04.041

Eickhoff SB, Stephan KE, Mohlberg H, Grefkes C, Fink GR, Amunts K, Zilles K. 2005. A new SPM toolbox for combining probabilistic cytoarchitectonic maps and functional imaging data. Neuroimage 25:1325–1335. doi:10.1016/j.neuroimage.2004.12.034

Fischer H, Wright CI, Whalen PJ, McInerney SC, Shin LM, Rauch SL. 2003. Brain habituation during repeated exposure to fearful and neutral faces: A functional MRI study. Brain Res Bull 59:387–392. doi:10.1016/S0361-9230(02)00940-1

Frässle S, Sommer J, Jansen A, Naber M, Einhäuser W. 2014. Binocular rivalry: Frontal activity relates to introspection and action but not to perception. J Neurosci 34:1738–1747. doi:10.1523/JNEUROSCI.4403-13.2014

Gayet S, Paffen CLE, Belopolsky A V., Theeuwes J, Van der Stigchel S. 2016. Visual input signaling threat gains preferential access to awareness in a breaking continuous flash suppression paradigm. Cognition. doi:10.1016/j.cognition.2016.01.009

Goeleven E, De Raedt R, Leyman L, Verschuere B. 2008. The Karolinska directed emotional faces: A validation study. Cogn Emot 22:1094–1118. doi:10.1080/02699930701626582

Gomes N, Silva S, Silva CF, Soares SC. 2017. Beware the serpent: the advantage of ecologically-relevant stimuli in accessing visual awareness. Evol Hum Behav 38:227–234. doi:10.1016/j.evolhumbehav.2016.10.004

Goossens L, Kukolja J, Onur OA, Fink GR, Maier W, Griez E, Schruers K, Hurlemann R. 2009. Selective processing of social stimuli in the superficial amygdala. Hum Brain Mapp 30:3332–3338. doi:10.1002/hbm.20755

Heekeren HR, Marrett S, Bandettini PA, Ungerleider LG. 2004. A general mechanism for perceptual decision-making in the human brain. Nature 431:859–862. doi:10.1038/nature02966

Heekeren HR, Marrett S, Ungerleider LG. 2008. The neural systems that mediate human perceptual decision making. Nat Rev Neurosci 9:467–479. doi:10.1038/nrn2374

Jiang Y, Costello P, He S. 2007. Processing of invisible stimuli: Advantage of upright faces and recognizable words in overcoming interocular suppression. Psychol Sci 18:349–355. doi:10.1111/j.1467-9280.2007.01902.x

Jiang Y, He S. 2006. Cortical Responses to Invisible Faces: Dissociating Subsystems for Facial-Information Processing. Curr Biol 16:2023–2029. doi:10.1016/j.cub.2006.08.084

Jiang Y, Shannon RW, Vizueta N, Bernat EM, Patrick CJ, He S. 2009. Dynamics of processing invisible faces in the brain: Automatic neural encoding of facial expression information. Neuroimage 44:1171–1177. doi:10.1016/j.neuroimage.2008.09.038

Kesler-West ML, Andersen AH, Smith CD, Avison MJ, Davis CE, Kryscio RJ, Blonder LX. 2001. Neural substrates of facial emotion processing using fMRI. Cogn Brain Res 11:213– 226. doi:10.1016/S0926-6410(00)00073-2

Knapen T, Brascamp J, Pearson J, van Ee R, Blake R. 2011. The role of frontal and parietal brain areas in bistable perception. J Neurosci 31:10293–10301. doi:10.1523/JNEUROSCI.1727-11.2011

Koller K, Rafal RD, Platt A, Mitchell ND. 2019. Orienting toward threat: Contributions of a subcortical pathway transmitting retinal afferents to the amygdala via the superior colliculus and pulvinar. Neuropsychologia 128:78–86. doi:10.1016/J.NEUROPSYCHOLOGIA.2018.01.027

Ledoux J. 1998. The Emotional Brain: The Mysterious Underpinnings of Emotional Life - Joseph Ledoux - Google Books. Simon and Schuster.

Lerche V, Bucher A, Voss A. 2019. Processing Emotional Expressions Under Fear of Rejection: Findings From Diffusion Model Analyses. Emotion 21:184. doi:10.1037/emo0000691

Lin H, Müller-Bardorff M, Gathmann B, Brieke J, Mothes-Lasch M, Bruchmann M, Miltner WHR, Straube T. 2020. Stimulus arousal drives amygdalar responses to emotional expressions across sensory modalities. Sci Rep 10:1–12. doi:10.1038/s41598-020-58839-1

Lufityanto G, Donkin C, Pearson J. 2016. Measuring Intuition: Nonconscious Emotional Information Boosts Decision Accuracy and Confidence. Psychol Sci 27:622–634. doi:10.1177/0956797616629403

Lumer ED, Friston KJ, Rees G. 1998. Neural correlates of perceptual rivalry in the human brain. Science (80-) 280:1930–1934. doi:10.1126/science.280.5371.1930

Lumer ED, Rees G. 1999. Covariation of activity in visual and prefrontal cortex associated with subjective visual perception. Proc Natl Acad Sci USA 96:1669–1673. doi:10.1073/pnas.96.4.1669

McFadyen J, Mermillod M, Mattingley JB, Halász V, Garrido MI. 2017. A Rapid Subcortical Amygdala Route for Faces Irrespective of Spatial Frequency and Emotion. J Neurosci 37:3864–3874. doi:10.1523/JNEUROSCI.3525-16.2017

McFadyen J, Smout C, Tsuchiya N, Mattingley JB, Garrido MI. 2019. SURPRISING THREATS ACCELERATE EVIDENCE ACCUMULATION FOR CONSCIOUS PERCEPTION. bioRxiv.

Melloni L, Schwiedrzik CM, Müller N, Rodriguez E, Singer W. 2011. Expectations change the signatures and timing of electrophysiological correlates of perceptual awareness. J Neurosci 31:1386–1396. doi:10.1523/JNEUROSCI.4570-10.2011

Menon V, Uddin LQ. 2010. Saliency, switching, attention and control: a network model of insula function. Brain Struct Funct 31:1386–1396. doi:10.1007/s00429-010-0262-0

Olszanowski M, Pochwatko G, Kuklinski K, Scibor-Rylski M, Lewinski P, Ohme RK. 2014. Warsaw set of emotional facial expression pictures: A validation study of facial display photographs. Front Psychol 5:1516. doi:10.3389/fpsyg.2014.01516

Ottaviani C, Cevolani D, Nucifora V, Borlimi R, Agati R, Leonardi M, De Plato G, Brighetti G. 2012. Amygdala responses to masked and low spatial frequency fearful faces: A preliminary fMRI study in panic disorder. Psychiatry Res - Neuroimaging 203:159–165. doi:10.1016/j.pscychresns.2011.12.010

Pasley BN, Mayes LC, Schultz RT. 2004. Subcortical discrimination of unperceived objects during binocular rivalry. Neuron 42:163–172. doi:10.1016/S0896-6273(04)00155-2

Pessoa L, Adolphs R. 2010. Emotion processing and the amygdala: From a “low road” to “many roads” of evaluating biological significance. Nat Rev Neurosci 11:773–782. doi:10.1038/nrn2920

Philiastides MG, Auksztulewicz R, Heekeren HR, Blankenburg F. 2011. Causal role of dorsolateral prefrontal cortex in human perceptual decision making. Curr Biol 21:980–983. doi:10.1016/j.cub.2011.04.034

Pinto Y, van Gaal S, de Lange FP, Lamme VAF, Seth AK. 2015. Expectations accelerate entry of visual stimuli into awareness. J Vis 15:13–13. doi:10.1167/15.8.13

Pleger B, Ruff CC, Blankenburg F, Bestmann S, Wiech K, Stephan KE, Capilla A, Friston KJ, Dolan RJ. 2006. Neural coding of tactile decisions in the human prefrontal cortex. J Neurosci 26:12596–12601. doi:10.1523/JNEUROSCI.4275-06.2006

Ploran EJ, Nelson SM, Velanova K, Donaldson DI, Petersen SE, Wheeler ME. 2007. Evidence accumulation and the moment of recognition: Dissociating perceptual recognition processes using fMRI. J Neurosci 27:11912–11924. doi:10.1523/JNEUROSCI.3522-07.2007

Prins N, Kingdom FA. 2009. Palamedes: Matlab routines for analyzing psychophysical data.

Ratcliff R. 1978. A theory of memory retrieval. Psychol Rev 85:59. doi:10.1037/0033-295X.85.2.59

Reinders AATS, Den Boer JA, Büchel C. 2005. The robustness of perception. Eur J Neurosci 22:524–530. doi:10.1111/j.1460-9568.2005.04212.x

Reinders AATS, Gläscher J, de Jong JR, Willemsen ATM, den Boer JA, Büchel C. 2006. Detecting fearful and neutral faces: BOLD latency differences in amygdala-hippocampal junction. Neuroimage 33:805–814. doi:10.1016/j.neuroimage.2006.06.052

Rounis E, Maniscalco B, Rothwell JC, Passingham RE, Lau H. 2010. Theta-burst transcranial magnetic stimulation to the prefrontal cortex impairs metacognitive visual awareness. Cogn Neurosci 1:165–175. doi:10.1080/17588921003632529

Salomon R, Ronchi R, Dönz J, Bello-Ruiz J, Herbelin B, Faivre N, Schaller K, Blanke O. 2018. Insula mediates heartbeat related effects on visual consciousness. Cortex 101:87–95. doi:10.1016/j.cortex.2018.01.005

Salomon R, Ronchi R, Dönz J, Bello-Ruiz J, Herbelin B, Martet R, Faivre N, Schaller K, Blanke O. 2016. The insula mediates access to awareness of visual stimuli presented synchronously to the heartbeat. J Neurosci 36:5115–5127. doi:10.1523/JNEUROSCI.4262-15.2016

Sander D, Grafman J, Zalla T. 2003. The Human Amygdala: An Evolved System for Relevance Detection. Rev Neurosci 14:303–316. doi:10.1515/REVNEURO.2003.14.4.303

Santos A, Mier D, Kirsch P, Meyer-Lindenberg A. 2011. Evidence for a general face salience signal in human amygdala. Neuroimage 54:3111–3116. doi:10.1016/j.neuroimage.2010.11.024

Sladky R, Geissberger N, Pfabigan DM, Kraus C, Tik M, Woletz M, Paul K, Vanicek T, Auer B, Kranz GS, Lamm C, Lanzenberger R, Windischberger C. 2018. Unsmoothed functional MRI of the human amygdala and bed nucleus of the stria terminalis during processing of emotional faces. Neuroimage 168:383–391. doi:10.1016/j.neuroimage.2016.12.024

Stein T, Hebart MN, Sterzer P. 2011. Breaking continuous flash suppression: A new measure of unconscious processing during interocular suppression? Front Hum Neurosci 5:167. doi:10.3389/fnhum.2011.00167

Stein T, Seymour K, Hebart MN, Sterzer P. 2014. Rapid Fear Detection Relies on High Spatial Frequencies. Psychol Sci 25:566–574. doi:10.1177/0956797613512509

Sterzer P, Haynes JD, Rees G. 2008. Fine-scale activity patterns in high-level visual areas encode the category of invisible objects. J Vis 8:10–10. doi:10.1167/8.15.10

Summerfield C, Egner T, Greene M, Koechlin E, Mangels J, Hirsch J. 2006. Predictive codes for forthcoming perception in the frontal cortex. Science (80-) 314:1311–1314. doi:10.1126/science.1132028

Tamietto M, de Gelder B. 2010. Neural bases of the non-conscious perception of emotional signals. Nat Rev Neurosci 11:697–709. doi:10.1038/NRN2889

Tipples J. 2015. Rapid Temporal Accumulation in Spider Fear: Evidence From Hierarchical Drift Diffusion Modelling. Emotion 15:742. doi:10.1037/emo0000079

Tottenham N, Tanaka JW, Leon AC, McCarry T, Nurse M, Hare TA, Marcus DJ, Westerlund A, Casey BJ, Nelson C. 2009. The NimStim set of facial expressions: Judgments from untrained research participants. Psychiatry Res 168:242–249. doi:10.1016/j.psychres.2008.05.006

Troiani V, Schultz RT. 2013. Amygdala, pulvinar, and inferior parietal cortex contribute to early processing of faces without awareness. Front Hum Neurosci 168:242–249. doi:10.3389/fnhum.2013.00241

Tsuchiya N, Koch C. 2005. Continuous flash suppression reduces negative afterimages. Nat Neurosci 8:1096–1101. doi:10.1038/nn1500

van der Schalk J, Hawk ST, Fischer AH, Doosje B. 2011. Moving Faces, Looking Places: Validation of the Amsterdam Dynamic Facial Expression Set (ADFES). Emotion 11:907. doi:10.1037/a0023853

Vizueta N, Patrick CJ, Jiang Y, Thomas KM, He S. 2012. Dispositional fear, negative affectivity, and neuroimaging response to visually suppressed emotional faces. Neuroimage 59:761– 771. doi:10.1016/j.neuroimage.2011.07.015

Voss A, Voss J. 2007. Fast-dm: A free program for efficient diffusion model analysis. Behav Res Methods 39:767–775. doi:10.3758/BF03192967

Wang C, Kang-Park MH, Wilson WA, Moore SD. 2002. Properties of the pathways from the lateral amygdal nucleus to basolateral nucleus and amygdalostriatal transition area. J Neurophysiol 87:2593–2601. doi:10.1152/jn.2002.87.5.2593

Willenbockel V, Sadr J, Fiset D, Horne G, Gosselin F, Tanaka J. 2010. The SHINE toolbox for controlling low-level image properties. J Vis. doi:10.1167/10.7.653

Williams MA, Morris AP, McGlone F, Abbott DF, Mattingley JB. 2004. Amygdala Responses to Fearful and Happy Facial Expressions under Conditions of Binocular Suppression. J Neurosci 24:2898–2904. doi:10.1523/JNEUROSCI.4977-03.2004

Yang E, Zald DH, Blake R. 2007. Fearful Expressions Gain Preferential Access to Awareness During Continuous Flash Suppression. Emotion 7:882. doi:10.1037/1528-3542.7.4.882

