## Supplementary Material for "Neural and computational processes of accelerated perceptual awareness and decisions: A 7T fMRI Study"

### SUPPLIMENTARY MATERIALS

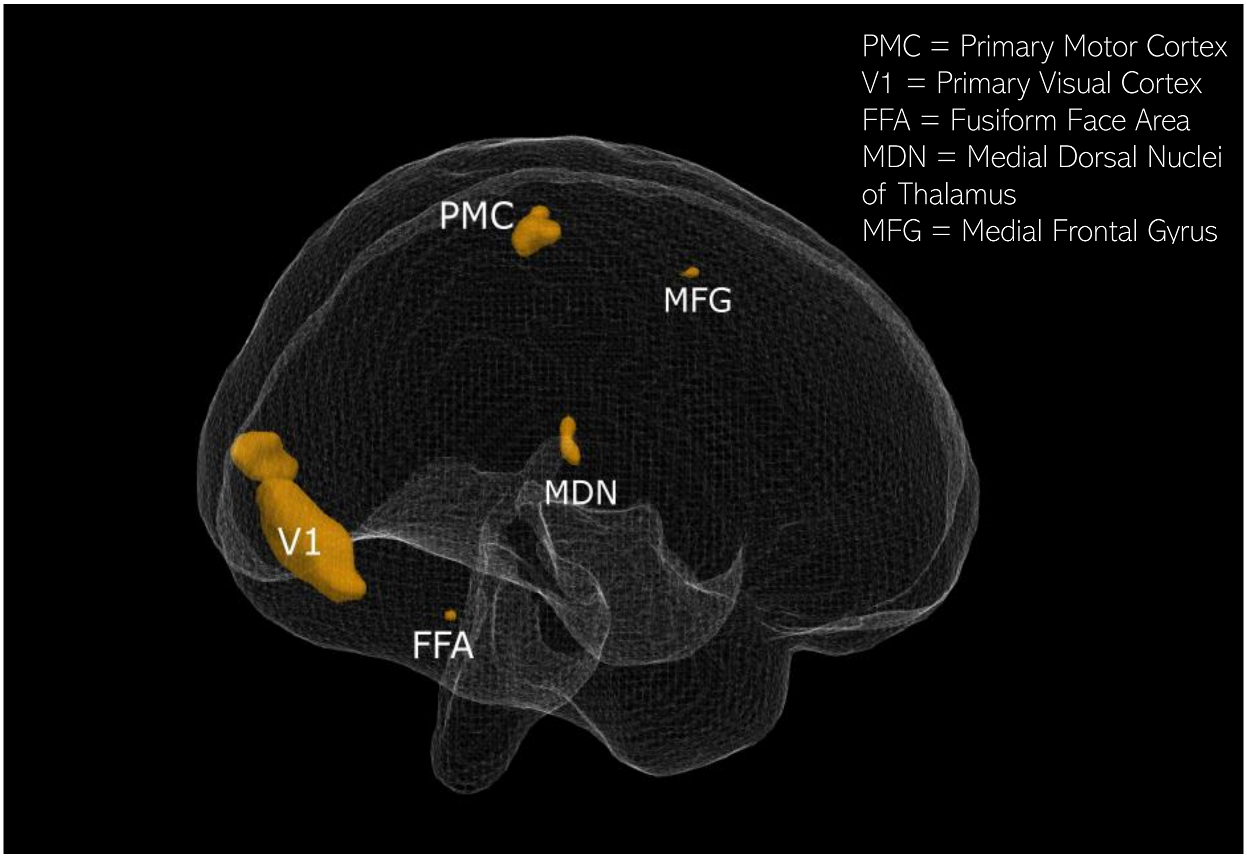


*Figure S1. Brain activity in conditions.* There were no brain activity differences between each of the four conditions; expected fear, expected neutral, unexpected fear and unexpected neutral. Each condition had activity in the brain regions highlighted above. This is an image of activity in just the expected fear condition as an example.

*Table S1: Brain activity correlates with faster reaction time*

| X | Y | Z | REGION | SIDE |
|---|---|---|---|---|
| 50 | -12 | 16 | Rolandic Operculum | Right |
| 39 | -16 | 16 | Insula Lobe | Right |
| -45 | -58 | -16 | Fusiform Gyrus | Left |
| 59 | 4 | 22 | Precentral Gyrus | Right |
| 44 | -3 | -12 | Superior Temporal Gyrus | Right |
| -44 | -21 | 14 | Rolandic Operculum | Left |
| -63 | -9 | 22 | Postcentral Gyrus | Left |
| -51 | -27 | 16 | IPL (SupraMarginal Gyrus) | Right |
| 60 | -51 | -9 | Middle Temporal Gyrus | Right |
| -42 | -24 | 37 | Postcentral Gyrus | Left |
| 17 | -82 | 46 | Cuneus | Right |
| 3 | -78 | 29 | Cuneus | Left |
| -11 | -84 | 40 | Superior Occipital Gyrus | Left |
| -54 | 38 | 5 | IFG (p. Triangularis) | Left |
| -45 | 4 | -12 | Superior Temporal Gyrus | Left |
| 60 | -9 | -4 | Superior Temporal Gyrus | Right |
| -41 | -12 | 11 | Insula Lobe | Left |
| -60 | -28 | 26 | IPL (SupraMarginal Gyrus) | Left |
| -50 | -52 | 5 | Middle Temporal Gyrus | Left |
| 33 | -16 | 47 | Precentral Gyrus | Right |
| -42 | -13 | 41 | Postcentral Gyrus | Left |
| -24 | 7 | -19 | Olfactory cortex | Left |
| 29 | 22 | -9 | Insula Lobe | Right |
| 8 | -75 | 2 | Lingual Gyrus | Right |
| 71 | -13 | 17 | Postcentral Gyrus | Right |
| 18 | -28 | -4 | Visual Thalamus | Right |
| 42 | -75 | 31 | Middle Occipital Gyrus | Right |
| -60 | 2 | 35 | Precentral Gyrus | Left |
| 50 | -45 | -12 | Middle Temporal Gyrus | Right |
| -6 | -48 | 47 | Precuneus | Left |
| 54 | 44 | 2 | IFG (p. Triangularis) | Right |
| -48 | -40 | 2 | Middle Temporal Gyrus | Left |
| 3 | -81 | 14 | Calcarine Gyrus | Left |
| 27 | -6 | -13 | Amygdala | Right |
| -26 | -84 | 41 | Middle Occipital Gyrus | Left |
| 39 | 34 | -6 | IFG (p. Orbitalis) | Right |
| -41 | 20 | -21 | Temporal Pole | Left |

*IPL = Inferior Parietal Lobule, IFG = Inferior Frontal Gyrus*

*Table S2: Brain activity correlates with increased drift-rate*

| X | Y | Z | REGION | SIDE |
|---|---|---|---|---|
| -38 | -45 | 49 | Inferior Parietal Lobule | L |
| -23 | -39 | -10 | ParaHippocampal Gyrus | L |
| -36 | -40 | -13 | Inferior Temporal Gyrus | L |
| -44 | -66 | -1 | Middle Occipital Gyrus | L |
| 51 | -42 | 14 | Superior Temporal Gyrus | R |
| 48 | -45 | 25 | SupraMarginal Gyrus | R |
| 45 | 19 | 35 | IFG (p. Opercularis) | R |
| 44 | 26 | 31 | IFG (p. Triangularis) | R |
| 45 | 22 | 46 | Middle Frontal Gyrus | R |
| 53 | -46 | 46 | Inferior Parietal Lobule | R |
| 23 | -34 | -13 | ParaHippocampal Gyrus | R |
| -24 | -88 | -18 | Lingual Gyrus | L |
| 51 | -63 | 40 | Angular Gyrus | R |
| -3 | -7 | 34 | MCC | L |
| 26 | -21 | -3 | Thalamus: Parietal | |
| -65 | -46 | 14 | Superior Temporal Gyrus | L |
| -39 | -19 | 8 | Heschls Gyrus | L |
| -15 | -73 | 14 | Calcarine Gyrus | L |
| -27 | -63 | 53 | Superior Parietal Lobule | L |
| 14 | 17 | 46 | Superior Frontal Gyrus | R |
| -24 | -40 | -33 | Cerebelum (IV-V) | L |
| -29 | -19 | -16 | Hippocampus | L |
| 26 | -55 | 22 | Precuneus | R |
| 29 | -67 | 25 | Superior Occipital Gyrus | R |
| -15 | -25 | 1 | Thalamus | L |
| 39 | -72 | 31 | Middle Occipital Gyrus | R |
| -36 | 5 | 52 | Middle Frontal Gyrus | L |
| -24 | -84 | 32 | Superior Occipital Gyrus | L |
| -56 | -64 | 20 | Middle Temporal Gyrus | L |
| -39 | -30 | 4 | Superior Temporal Gyrus | L |
| -6 | -63 | 25 | Precuneus | L |
| -14 | -66 | 25 | Cuneus | L |
| -12 | -63 | 16 | Calcarine Gyrus | L |
| 57 | -46 | -12 | Inferior Temporal Gyrus | R |
| 20 | 25 | 46 | Superior Frontal Gyrus | R |
| 6 | -22 | 17 | Thalamus: Temporal | |
| 42 | -16 | 7 | Heschls Gyrus | R |
| -14 | -49 | 7 | Calcarine Gyrus | L |
| -8 | -45 | 13 | PCC | L |

| | | | | |
|---|---|---|---|---|
| -36 | 5 | 44 | Precentral Gyrus | L |
| 18 | -25 | 47 | MCC | R |
| 33 | -57 | 49 | Inferior Parietal Lobule | R |
| 8 | -90 | -1 | Calcarine Gyrus | R |
| 71 | -42 | 1 | Middle Temporal Gyrus | R |
| 21 | -13 | -21 | ParaHippocampal Gyrus | R |
| -8 | -87 | 25 | Cuneus | L |
| 59 | 4 | -18 | Medial Temporal Pole | R |
| 11 | -82 | 8 | Calcarine Gyrus | R |
| -17 | -69 | 43 | Superior Parietal Lobule | L |
| -20 | -22 | 10 | Thalamus | L |
| 12 | -49 | 23 | Precuneus | R |
| -12 | -73 | -22 | Cerebelum (VI) | L |
| -56 | -40 | -4 | Middle Temporal Gyrus | L |
| -63 | -46 | 25 | SupraMarginal Gyrus | L |
| -23 | -12 | -16 | Amygdala | L |

*IFG = Inferior Frontal Gyrus, MCC = mid cingulate cortex, PCC = posterior cingulate cortex*
